# When Wavelengths Collide: Bias in Cell Abundance Measurements due to Expressed Fluorescent Proteins

**DOI:** 10.1101/037010

**Authors:** Ariel Hecht, Drew Endy, Marc Salit, Matthew S. Munson

## Abstract

The abundance of bacteria in liquid culture is commonly inferred by measuring optical density at 600 nm. Red fluorescent proteins (RFP) can strongly absorb light at 600 nm. Increasing RFP expression can falsely inflate apparent cell density and lead to underestimations of mean per-cell fluorescence by up to 10%. Measuring optical density at 700 nm would allow estimation of cell abundance unaffected by the presence of nearly all fluorescent proteins.

---

A common method for estimating the abundance of bacteria in a liquid culture is to measure the optical density (OD) of the culture at a wavelength of 600 nm (OD_600_)^1,2^. The OD of the sample is defined as the negative log of the attenuation of the incident light:

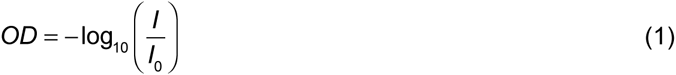

where *I_0_* is the intensity of incident light shined at the sample, and *I* is the intensity of light that reaches the photodetector after passing through the sample.

When estimating cell density in liquid culture, it is commonly assumed that the dominant source of the attenuation of incident light is scattering by cells.^3^ If cells express fluorescent proteins that absorb light at the wavelength used for measuring OD then this absorption can also attenuate incident light (Figure 1). Such effects have been previously reported for microorganisms that express naturally-occurring pigments.^4-6^ Here, we quantify the extent of such error due to heterologous expression of fluorescent proteins.

**Figure 1.**
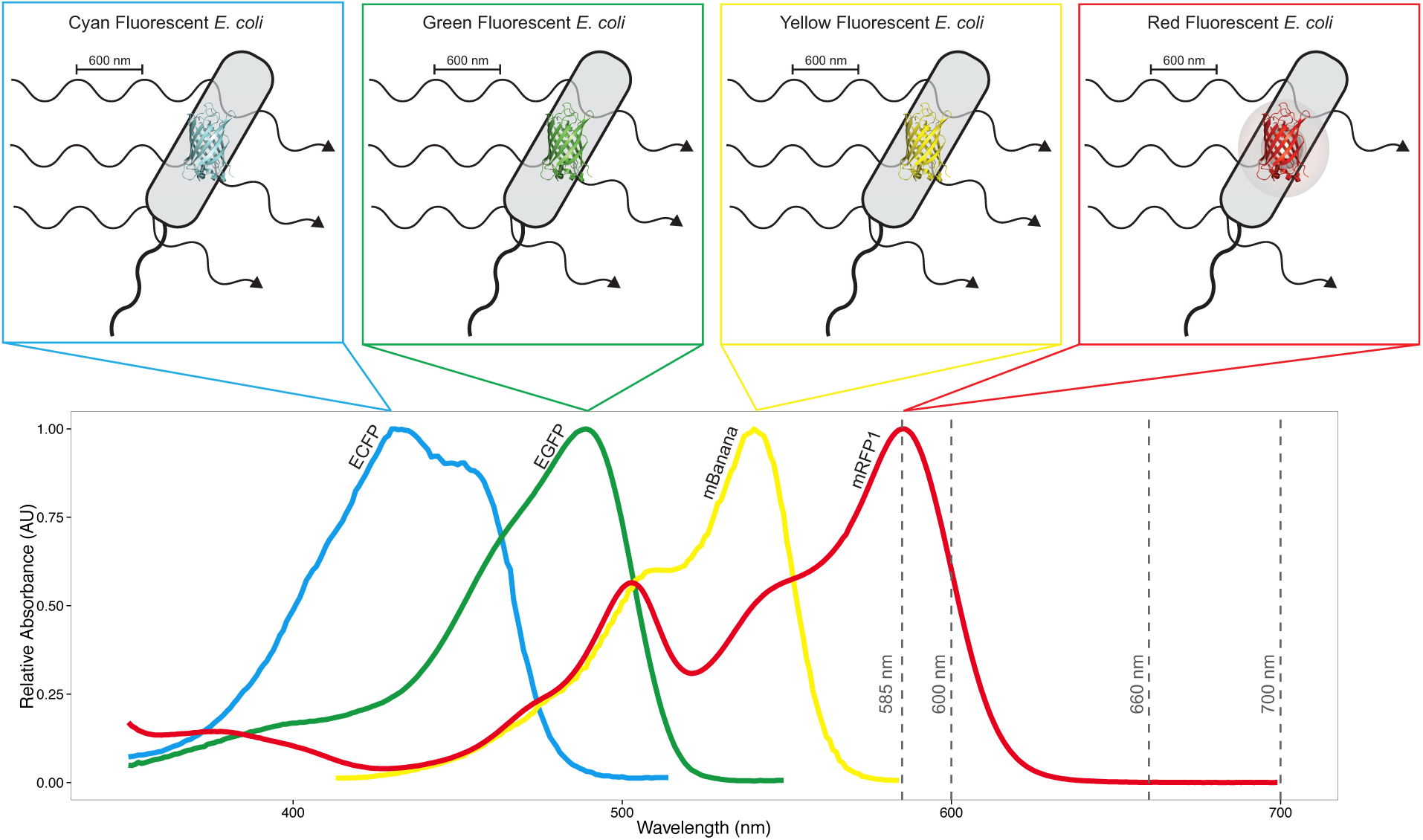
Absorption biases estimates of cell abundance from measurements of OD_600_ in cells expressing red fluorescent proteins. When OD_600_ is measured for *Escherichia coli (E. coli)* that do not express fluorescent proteins, the dominant source of attenuation of incident light is scattering. When OD600 is measured for *E. coli* that express red fluorescent proteins (RFP), incident light is attenuated both by scattering and by absorption via the RFPs, which in turn biases the calculation of per-cell fluorescence. This additional source of attenuation biases estimates of cell abundance from measurements of OD_600_. Absorption spectra of fluorescent proteins from four parts of the visible color spectrum are shown: cyan (ECFP), green (EGFP), yellow (mBanana) and red (mRFP1).^13^ Of these fluorescent proteins, only mRFP1 has significant absorbance at 600 nm. The four dashed vertical lines are at the four wavelengths at which OD was measured for this study: 585 nm, 600 nm, 660 nm, and 700 nm.

Fluorescent proteins are often incorporated as reporters in synthetic genetic constructs, where their expression can be quantified as a proxy for the activity of the construct under study. The bias in the estimation of cell abundance from the measurement of OD is of particular concern when the objective is to measure per-cell fluorescence, which is then utilized to quantitatively compare expression under different experimental conditions. These measurements are commonly used in synthetic biology and genetic engineering for investigating the impact of genetic composition on expression, and balancing expression levels for different elements in a pathway or system.^7-12^

Avoiding biased estimates of cell abundance arising from non-scattering attenuation can be accomplished by measuring OD at a wavelength that is not absorbed by the reporter, or by selecting a reporter that does not absorb at the wavelength used to measure OD for cell density estimation. When a single reporter is used, the selection of an alternative reporter is straightforward. However, when multiple reporters are required avoiding the use of RFP can be challenging, while measuring OD at an alternative wavelength remains straightforward.

Frequently used RFPs that are likely to introduce bias include mRFP1 (Figure 1); mCherry, which has an absorption peak at 587 nm;^14^ and mKate2, which has an absorption peak at 588 nm.^15^ Despite evidence showing that estimates of cell abundance from measurements of OD can be biased by absorption,^4-6^ and the simplicity with which the bias can be avoided, the synthetic biology community does not yet seem to have adopted alternatives that avoid such errors (Table 1).

**Table 1.** Number of hits returned for a full-text Google Scholar search of the combination of the terms for the fluorescent protein mCherry, the bacteria Escherichia coli and wavelengths used for OD measurements.

| Search Terms | Hits |
| --- | --- |
| mCherry AND “Escherichia coli” AND “OD 600” | 1010 |
| mCherry AND “Escherichia coli” AND “OD 660” | 46 |
| mCherry AND “Escherichia coli” AND “OD 700” | 1 |

One example of such biasing can be found in Mutalik, *et al.^9^* In this paper, the authors investigated the impact of interactions between genetic elements by characterizing the per-cell fluorescence from a library of promoters and 5'-UTRs driving the expression of both sfGFP and mRFP1. They concluded that approximately 14% of the variation in fluorescence is attributed to interactions with the reporter gene. However, at least some of this variation seems attributable to the bias from measuring OD_600_ in RFP-expressing cells.

In another example, both GFP and RFP are measured simultaneously in liquid culture to correlate the impact of the burden imposed by synthetic constructs (mCherry) on the expression capacity of the cell (sfGFP). When these authors normalize fluorescence by the cell density estimate from OD_600_, a bias is introduced in some of the reported values of the metabolic burden imposed by various constructs. This bias is unlikely to effect the conclusions of this method paper, but quantitative results of the method as reported are subject to systematic error. Normalizing fluorescence using OD_700_ would eliminate the bias and increase the quantitative accuracy of this method.

Here, we experimentally quantified the bias in estimates of cell abundance from measurements of OD using a set of *Escherichia coli (E. coli)* strains engineered to express mRFP1 (see Methods section).^10^ The fluorescence of each culture was measured at an excitation of 585 nm and an emission of 610 nm. The OD of each culture was measured at four wavelengths: 585 nm, 600 nm, 660 nm and 700 nm. The two shorter wavelengths are significantly absorbed by mRFP1, while the two longer wavelengths have negligible absorbance by this reporter (see Figure 1). We used OD_700_ as the benchmark of attenuation due to scattering to estimate the magnitude of the OD bias at the three shorter wavelengths.

To evaluate bias using OD_700_, we need to estimate the OD at other wavelengths, and consider the observed OD at any wavelength as the sum of two sources of attenuation (scattering and absorbance):

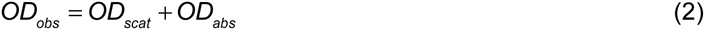

where *OD_obs_* is the observed OD, *OD_scat_* is the OD due to scattering by the cells, and *OD_abs_* is the OD bias arising from absorbance by expressed fluorescent proteins. To estimate *OD_scat_* at wavelengths between 500 nm and 800 nm for *E. coli*, we use the relationship described by Myers, *et al*. [6]:

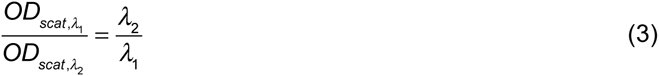

where *λ* is the wavelength of the incident light.

OD at each wavelength was plotted against fluorescence expression (Figure 2a). We observed a strong relationship between fluorescence and cell density - more cells resulted in higher total fluorescence. Using OD_700_ as an unbiased measure of scatter, we could account for the variation in cell density from culture to culture, and evaluated the OD bias at the other wavelengths. The absolute (Figure 2b) and relative (Figure 2c) bias were a function of wavelength.

**Figure 2.**
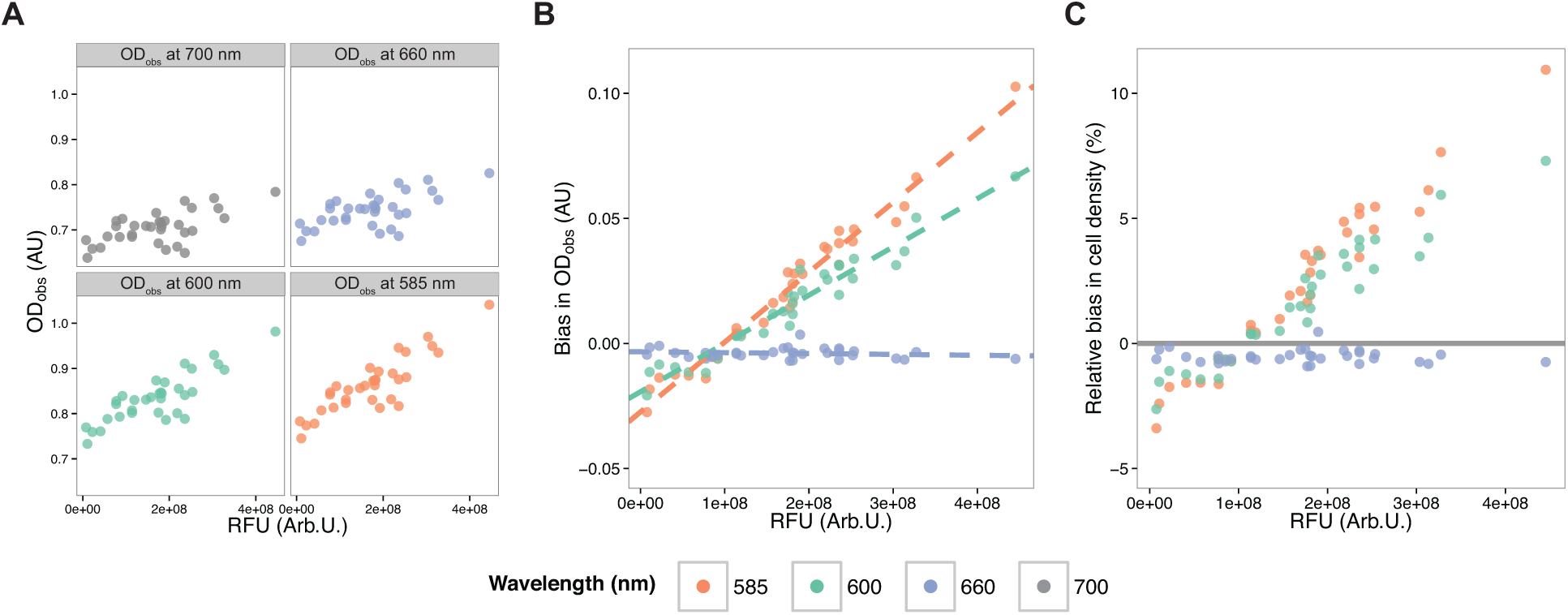
Estimates of cell abundance from measurements of OD at absorbed wavelengths are linearly biased by the absorption of incident light by fluorescent proteins. (A) *OD_obs_* of liquid cultures of cells expressing mRFPI measured at four wavelengths (585 nm, 600 nm, 660nm and 700nm) as a function of the measured fluorescence (excitation at 585 nm, emission at 610 nm). As the total fluorescence of the culture increased, the difference between *OD_obs_* at absorbing and non-absorbing wavelengths increased. (B) Bias in apparent cell abundance at 3 wavelengths, benchmarked against a non-absorbing wavelength, 700 nm. At absorbing wavelengths, the bias was linearly proportional to fluorescence. Dashed lines represent the linear fit of *OD_abs_* as a function of bulk fluorescence. The bias in estimating per-cell fluorescence was a linear function of total culture fluorescence when measured at absorbing wavelengths, with the impact greater as the wavelength approaches the mRFP1 absorbance peak. (C) Relative bias in *OD_obs_* due to the absorbance of incident light by mRFP1.

At the absorbing wavelengths (585 nm and 600 nm), we observed that *OD_absλ_* was linearly correlated with the total fluorescence of the culture, in agreement with the Beer-Lambert law (as shown by the dashed lines in Figure 2b). The non-absorbing “test” wavelength (660 nm) acted as a control, with *OD_abs, 660_* showing no relationship with the fluorescence of the culture. This validated our use of OD_700_ as a benchmark, and Eqs. (2) and (3).

At 585 nm, the relative bias in estimating cell abundance exceeded 10% for the highest-expressing cultures (Figure 2c). At 600 nm, the bias exceeded 7% for the highest-expressing cultures (Figure 2c), consistent with the absorbance spectrum of mRFP1 (Figure 1).

When the wavelength used to measure OD is absorbed by an expressed fluorescent protein, estimates of cell abundance from measurements of OD will be biased in proportion to the amount of mature fluorescent protein. Measurements of per-cell fluorescence will be similarly biased.

We also considered that a reciprocal bias (a bias in measuring fluorescence as a function of OD) may occur, which can arise from an “inner filter effect,”^16^ where both incident excitation radiation and fluorescent emission are attenuated. This effect can be corrected mathematically,^16^ or by diluting high OD samples prior to measurement. This does not appear to be a significant source of bias at OD_600_ less than 4,^17,18^ which is outside the range of measurements reported here and outside the sensitivity of most spectrometers.

We recommend that when estimating the per-cell fluorescence in a liquid culture OD should be measured at a wavelength that is not absorbed by any expressed fluorescent protein. Adopting the general practice of measuring OD at 700 nm, in place of the customary 600 nm, would allow unbiased estimates of cell abundance from measurements of OD of cells expressing nearly all visible-spectrum fluorescent proteins. As the practice of engineering cells becomes ever more precise there is a corresponding need for better characterization of the impact of genetic elements on gene expression. Adopting this simple change will improve the quality of data generated.

## Methods

We experimentally quantify the bias in estimates of cell abundance from measurements of OD using a set of *Escherichia coli (E. coli)* strains engineered to express mRFP1.^10^ These strains (Addgene kit# 1000000037) express mRFP1 under the control of a combinatorial set of constitutive Ptrc-variant promoters and bicistronic design (BCD) ribosome binding sites (RBS). Cell cultures were started by inoculations from frozen glycerol stocks into 500 μL of lysogeny broth (LB) containing 0.1 mmol L^-1^ (50 μg mL^-1^ ) kanamycin in 2 mL deep-well plates sealed with an AeraSeal gas-permeable microplate seal (E&K Scientific) and grown overnight (16 hours) at 37 °C in a Kuhner LT-X (Lab-Therm) incubator shaking at 460 rpm with 80 % humidity.^i^ Aliquots (150 μL) of the cultures were transferred to a CELLSTAR black, clear-bottom 96-well plate (Grenier, #M0562) and measured on a Molecular Devices SpectraMax i3 plate reader. The fluorescence of each culture was measured at an excitation of 585 nm and an emission of 610 nm. The OD of each culture was measured at four wavelengths: 585 nm, 600 nm, 660 nm and 700 nm.

## Footnotes

i Certain commercial equipment, instruments, or materials are identified in this report to specify adequately the experimental procedure. Such identification does not imply recommendation or endorsement by the National Institute of Standards and Technology, nor does it imply that the materials or equipment identified are necessarily the best available for the purpose

